# *Parp1* deletion rescues cerebellar hypotrophy in *xrcc1* mutant zebrafish

**DOI:** 10.1101/2024.11.25.625242

**Authors:** Svetlana A. Semenova, Deepthi Nammi, Grace A. Garrett, Gennady Margolin, Jennifer L. Sinclair, Reza Maroofian, Keith W. Caldecott, Harold A. Burgess

## Abstract

Defects in DNA single-strand break repair are associated with neurodevelopmental and neurodegenerative disorders. One such disorder is that resulting from mutations in *XRCC1*, a scaffold protein that plays a central role in DNA single-strand base repair. XRCC1 is recruited at sites of single-strand breaks by PARP1, a protein that detects and is activated by such breaks and is negatively regulated by XRCC1 to prevent excessive PARP binding and activity. Loss of XRCC1 leads to the toxic accumulation and activity of PARP1 at single-strand breaks leading to base excision repair defects, a mechanism that may underlie pathological changes in patients carrying deleterious *XRCC1* mutations. Here, we demonstrate that *xrcc1* knockdown impairs development of the cerebellar plate in zebrafish. In contrast, *parp1* knockdown alone does not significantly affect neural development, and instead rescues the cerebellar defects observed in *xrcc1* mutant larvae. These findings support the notion that PARP1 inhibition may be a viable therapeutic candidate in neurological disorders.

## Introduction

DNA single-strand breaks (SSB) are among the most common DNA lesions arising in cells, and are induced both by exposure to endogenous and exogenous (environmental) genotoxins, as a result of an intrinsic level of DNA instability, and as intermediates or byproducts of DNA metabolic processes ^1^. The threat posed by unrepaired SSBs to human health is exemplified by the existence of hereditary disorders in which genes encoding proteins that facilitate DNA single-strand break repair are mutated, and typically which are associated with neurodevelopmental dysfunction and/or neurodegeneration ^2^. One such disorder is spinocerebellar ataxia autosomal recessive 26 (SCAR26), which results from mutations in *XRCC1*; a gene encoding a scaffold protein that plays a central role in DNA single-strand base repair ^3,4^. PARP1, a protein that detects and is activated by SSBs, recruits XRCC1 to SSB sites where it negatively regulates PARP1 to prevent excessive binding and activity ^5–7^. The latter leads to excessive depletion of the NAD+ cofactor and metabolic dysfunction, and/or to toxic levels of poly(ADP-ribose)product. The importance of this relationship is illustrated by the observation that *Parp1* deletion suppresses the cerebellar defects and seizures in mice with *Xrcc1* deletion in brain ^4,8^. To what extent this relationship is reflected in other animal species is unclear, however. Here, we have examined the impact of *xrcc1* and/or *parp1* depletion in zebrafish, and we show that *parp1* knockdown rescues the cerebellar defects observed in *xrcc1* mutant larvae.

## Results

To characterize contributions of *xrcc1* to neural development we generated G0 *xrcc1* crispants. Co-injection of multiple highly efficient guide RNAs (gRNAs) can generate zebrafish G0 animals with bi-allelic mutations (crispants) that reproduce phenotypes in stable mutations ^9^. As a first step to generating *xrcc1* crispants, we injected a set of four guide RNAs that each targeted a different early exon of *xrcc1* to assess the cutting efficiency of each. gRNA-t1 and gRNA-t2 depleted the wildtype peak to less than 50% of the total product (Supp. Fig. 1). Thus, in subsequent experiments we co-injected gRNA-t1 and gRNA-t2 to generate *xrcc1* crispants. To minimize the possibility of off-target mutations, we also tested an independent set of guide RNAs that had no sequence overlap with gRNA-t1 or gRNA-t2, and identified two additional highly efficient gRNAs (gRNA-t5 and gRNA-t6 ; Supp. Fig. 1). We then focused on phenotypes that were present both in gRNA-t1/2 and gRNA-t5/6 crispants.

Xrcc1 crispants appeared normal during the first week of development, including inflating swimbladders at around 4 dpf (days post-fertilization) and beginning to swim freely. We measured balance, sensory responsiveness to auditory and visual cues, movement kinematics and sensorimotor integration, but did not detect any significant changes in behavior in 6 dpf larvae (Supp. Fig. 2). Thus, *xrcc1* knockout zebrafish show grossly normal development at early larval stages.

Loss of *xrcc1* is expected to lead to an accumulation of single-strand breaks, triggering expression of DNA-damage response genes such as p53 ^10,11^. We performed RNA-seq on crispants and controls from RNA isolated from head tissue to assess differentially expressed genes in crispants. In gRNA-t1/2 crispants, 125 genes were significantly changed and 842 in gRNA-t5/6 crispants. Of these 31 genes appeared in both lists, with the change occurring in the same direction, a highly significant convergence (Fisher Exact test p=1.9×10^−14^; Fig. 1A). All but three of these overlapping genes were upregulated, and unexpectedly, 25 were lens proteins (including 18 crystallins). Expression of *xrcc1* itself was significantly reduced, possibly reflecting nonsense-mediated decay of truncated transcripts. Importantly, p53 was elevated in both experiments, indicating that crispants reproduce a key *xrcc1* phenotype seen in mammalian models (Fig. 1B).

**Figure 1.**
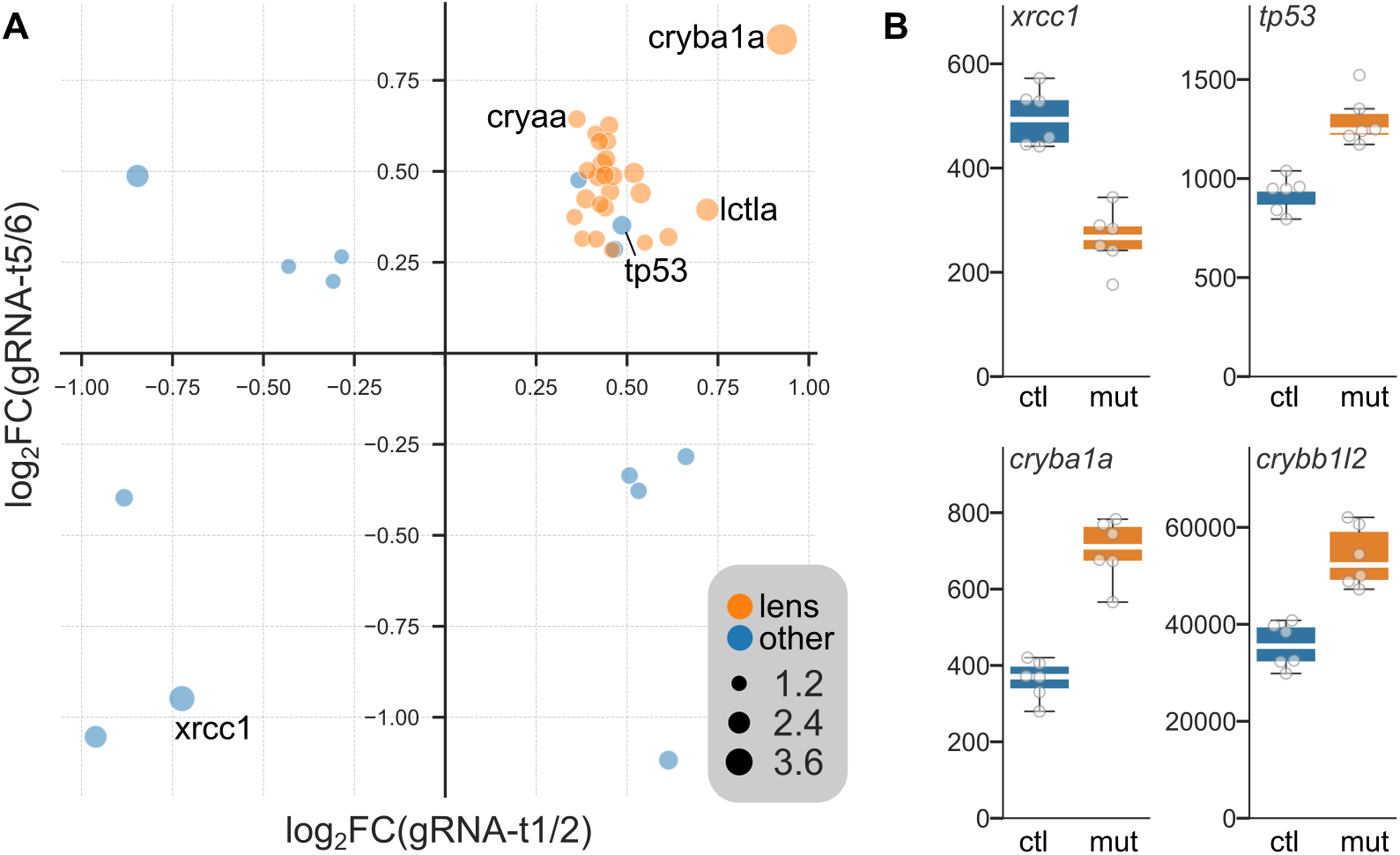
Differentially expressed genes in *xrcc1* crispants. A. Genes that were significantly changed in both gRNA-t1/2 and gRNA-t5/6 crispants. Axes show - 1×log_2_ (fold change) in crispants compared to controls. Dot size indicates log_10_(mean false discovery rate). Lens proteins are highlighted in orange. B. Normalized counts in controls (ctl) and crispants (mut) for selected genes that were differentially expressed in both experiments.

Next, we employed brain morphometry to assess whether crispants showed changes in brain architecture or composition ^12^. To do so, we co-injected gRNA-t1 and gRNA-t2 into triple transgenic fish in which all neurons were labeled with mCardinal (mCar), glutamatergic neurons labeled with GFP and GABAergic neurons labeled with RFP. After whole-brain confocal imaging of GFP, RFP and mCar in 6 dpf larvae, we performed elastic registration of each image stack to an annotated zebrafish brain atlas, allowing us to compare the fluorescence intensity in controls and mutants at each voxel, as well as in atlas-defined brain divisions ^12^. In addition, we used the inverse of the transformation matrix generated during registration to apply the atlas annotations to the original brain scans, allowing us to measure the volume of each brain, its divisions and individual voxels. Crispants showed no reproducible change in overall brain volume and no changes in either the mean fluorescence intensity or volume of any brain division (Supp. Fig. 3). However, when comparing maps of voxel volume (i.e., where individual voxels had to be dilated or compressed to match the reference brain), we noticed that there was a strong bilaterally symmetric localized reduction in voxel volume in part of the cerebellum in crispants (Fig. 2A). The affected region was in a lateral part of the cerebellar plate (LCeP), immediately above the cerebellar commissure that separates the cerebellum from the prepontine tegmentum. To determine whether this phenotype was reproducible, we made a mask from the affected region in gRNA-t1/2 crispants, and measured its volume in a second experiment using gRNA-t5/6. The same subregion of the cerebellar plate was significantly reduced (t-test, p = 0.0043; Fig. 2B), confirming that knock-down of *xrcc1* leads to a cerebellar phenotype in larval zebrafish.

**Figure 2.**
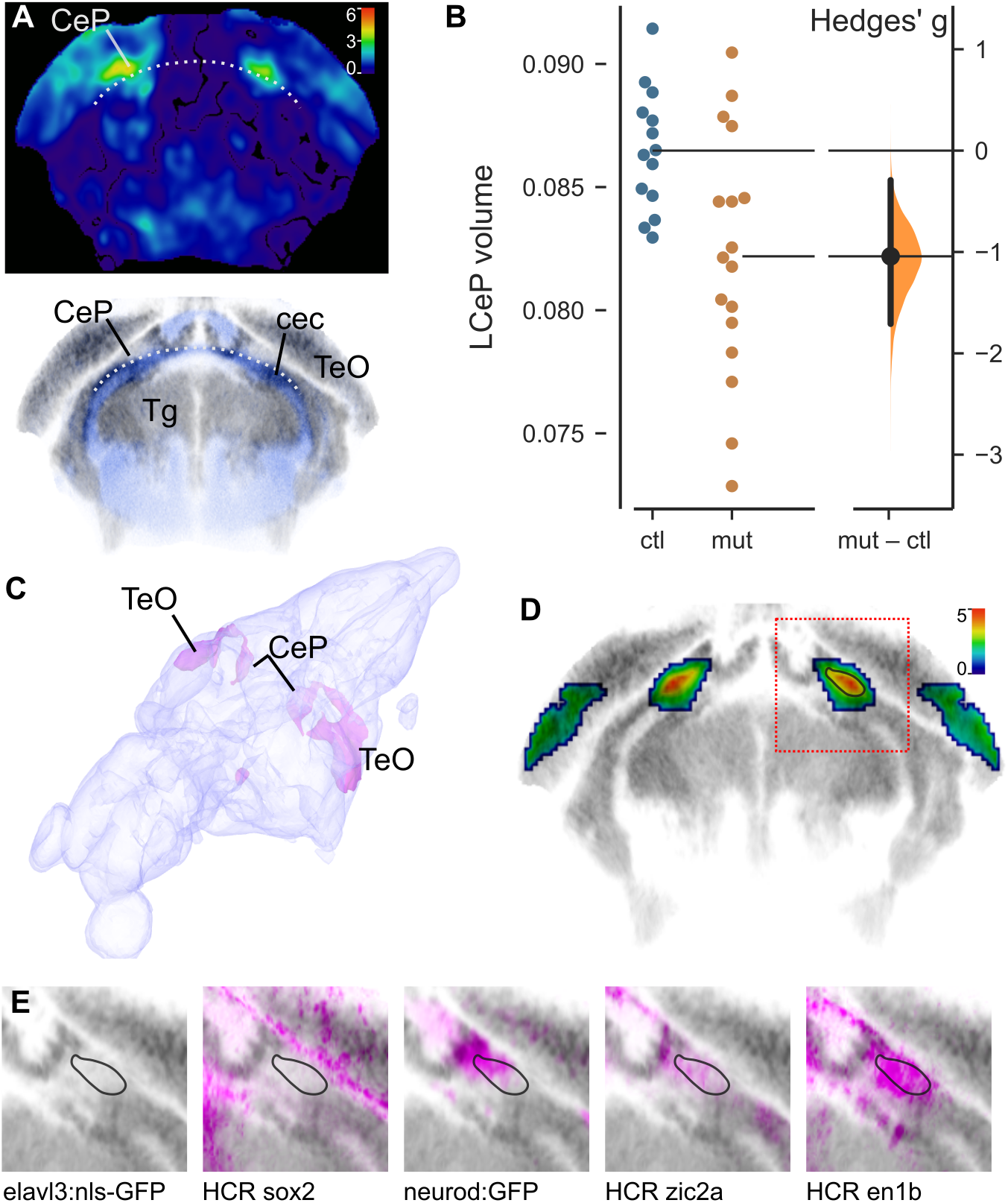
Reduced cerebellar plate volume in *xrcc1* crispants. A. Coronal section through the cerebellum of the voxelwise p-value map for voxel volume in controls versus mutants. Color bar indicates -1×log_10_(p-value). Lower panel: Same section from ZBB atlas showing cells (grey) and neuropil (blue). Dotted line marks the upper boundary of the cerebellar commissure (cec) that separates the cerebellar plate (CeP) from the prepontine tegmentum (Tg). At this level, CeP is positioned ventrally to the midbrain optic tectum (TeO). B. Volume (as percentage of total brain volume) of the affected part of the lateral cerebellar plate (LCeP) in second experiment using gRNAs-t5/6. N = 14 controls, 17 crispants. C. 3D projection from the Zebrafish Brain Browser superimposing on a brain model, the location of voxels that during brain registration, had to be inflated more in crispants than in controls (indicating areas that were smaller in the crispants). D. Coronal section (same plane as in A), through map of p-values for changes in voxel volume in *xrcc1* crispants derived from combining gRNA-t1/2 and gRNA-t5/6 experiments using ANOVA at each voxel. Color scale indicates -log_10_(p-value). p-values greater than 0.01 removed. Red square shows region shown in E. Black outline in right cerebellum corresponds to outlines in E. E. Characterization of the affected area by comparison to expression in transgenic lines elavl3:nls-GFP (marking neuronal somas) and neurod:GFP, and hybridization chain reaction (HCR) fluorescent in situ images for sox2, zic2a and en1b.

To more precisely localize the hypotrophic area, we performed a two-way ANOVA on each voxel in samples from the two experiments controlling for main effects of experiment and genotype, then filtered the resultant p-value map to exclude small clusters and voxels that were not bilaterally symmetric. In a 3D projection it was apparent that the area that was reduced in size in crispants extended dorsally from the LCeP and rostrally into the caudal aspect of the optic tectum (Fig. 2C). Using the Zebrafish Brain Browser (ZBB) atlas ^13^, we examined previously registered transgenic lines and in situ hybridization images to characterize the most strongly affected region (black outline in Fig. 2D). Intriguingly, this region was in an area containing a relatively low signal for mature neuronal somas (*elavl3:nls-GFP*, Fig. 2E), but that also lacked expression of a marker of neuronal progenitors (*sox2*). There was partial overlap with *neurod:GFP* and *zic2a*, markers of granule cells and granule cell progenitors respectively. However, the strongest overlap was with *en1b*, a gene that in mouse promotes the differentiation and survival of multiple cerebellar neuron subtypes ^14^.

In cultured mammalian cells lacking XRCC1, the poly(ADP-ribose) polymerase PARP1 becomes trapped on chromosomes during DNA base excision repair leading to repair failure and cell death. Accordingly, depletion or deletion of PARP1 can rescue DNA damage sensitivity in XRCC1 mutant cell lines ^4,6,8^. We therefore examined whether the cerebellar phenotype in *xrcc1* crispants was rescued if we simultaneously depleted *parp1*. We tested several gRNAs against *parp1* and identified four guides that were especially efficient at cutting their target sites (Supp. Fig. 4). We first checked for brain phenotypes in *parp1* crispants, again performing two experiments, each with two non-overlapping guide RNAs. We did not detect changes in the fluorescence intensity of the transgenic reporters, or size of any brain division, or local changes in brain volume in *parp1* crispants (Supp. Fig. 5). Importantly, the LCeP that was contracted in *xrcc1* crispants was unchanged in *parp1* crispants (Fig. 3A). Next, we tested the effect of simultaneously targeting *xrcc1* and *parp1*, compared to controls and to *xrcc1* alone. An ANOVA test confirmed that the volume of the LCeP was strongly affected by experimental condition (F_2,39_ = 5.98, p = 0.005). Post hoc comparisons confirmed that – as expected – the volume was smaller in *xrcc1* crispants than in controls. However, in double *xrcc1, parp1* crispants, the LCeP was significantly larger than in *xrcc1* crispants and in fact similar in size to controls, thus confirming that *parp1* deletion rescues cerebellar plate hypotrophy in *xrcc1* mutants (Fig. 3B).

**Figure 3.**
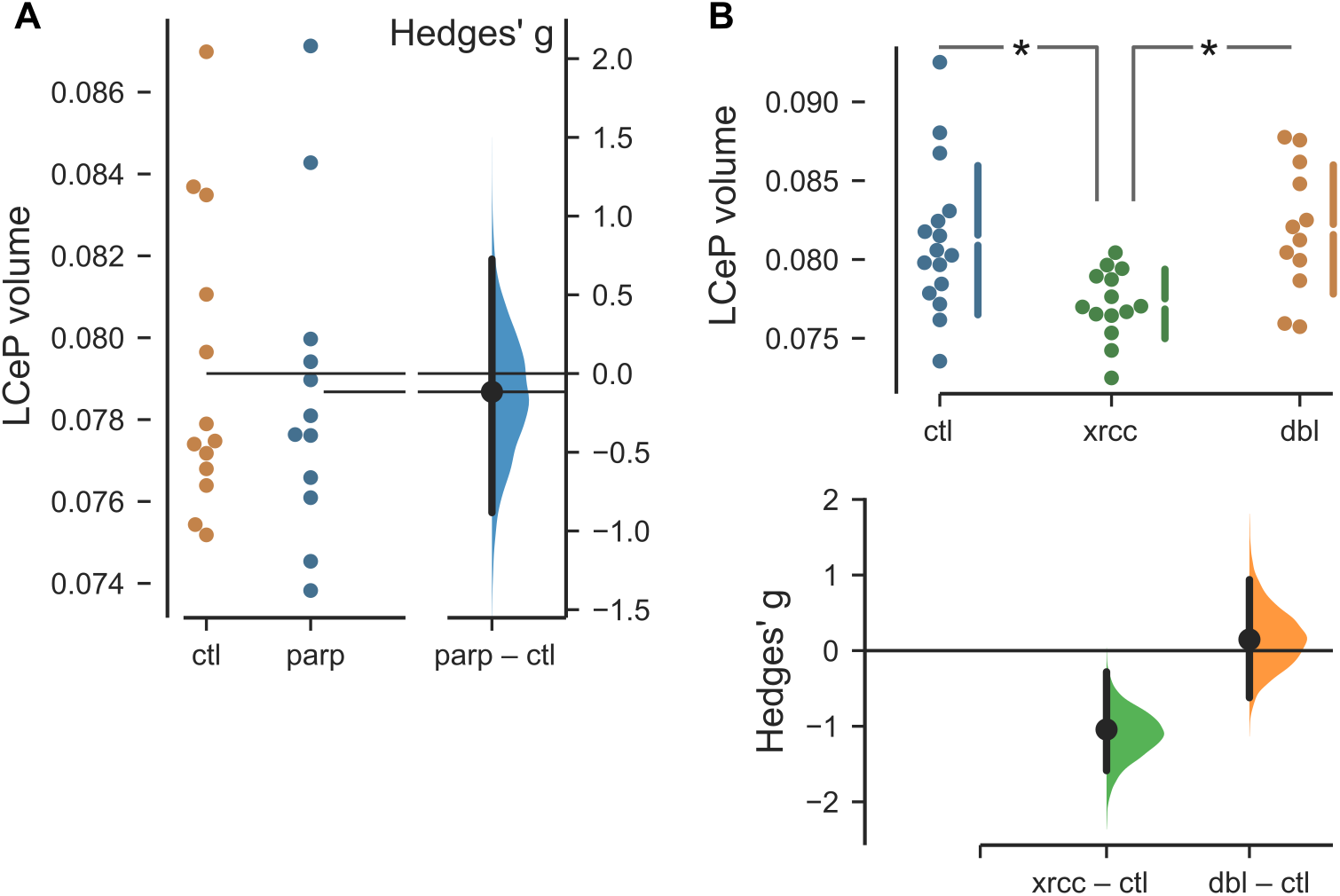
Rescue of *xrcc1* crispant cerebellar phenotype by *parp1* deletion. A.Volume of the lateral cerebellar plate (LCeP) in controls (ctl) and *parp1* crispants (parp). N=13,12. B.Volume of the LCeP in control larvae (ctl), *xrcc1* crispants (xrcc ; gRNA-t1/2) and *xrcc1, parp1* double crispants (dbl ; *xrcc1* gRNA-t1/2 and *parp1* gRNA-t6/8). N=16,14,12 respectively. * t-test p < 0.01

## Discussion

Defects in DNA single-strand break repair are associated with at least six neurodevelopmental and/or neurodegenerative disorders, one of which is caused by mutations in the central SSB repair scaffold protein, XRCC1. This disease, denoted spinocerebellar ataxia autosomal recessive 26 (SCAR26), is associated with progressive cerebellar degeneration and ataxia. Intriguingly, patient-derived cells from SCAR26 exhibit hyperactivity of the SSB sensor protein PARP1, a phenotype that is recapitulated in other human cell and mouse models of Xrcc1-defective disease. Here, we show that like Xrcc1-defective mice, *xrcc1* mutant zebrafish exhibit cerebellar defects, and that these defects are rescued by additional depletion of *parp1* ^4,8^. In particular, we show here that knockdown of *parp1* during zebrafish development rescues localized hypotrophy in a lateral part of the cerebellar plate in *xrcc1* mutant fish. This discovery is consistent with previous work showing that a key role of mammalian XRCC1 is to suppress toxic PARP1 hyperactivity during base excision repair ^5,6^. These data support the idea that inhibitors of Parp activity, which are currently employed in the clinic to treat certain types of breast and ovarian cancer, might potentially be repurposed for the treatment of Xrcc1-defective neurological disease.

An unexpected finding from RNA sequencing experiments was that a large number of lens proteins were upregulated in *xrcc1* crispants. Most of these were crystallins, many of which are related to heat-shock proteins and could be acting as stress-related chaperones. However, we also noted elevated expression of *lctla* (*lactase-like a*) and *calpain 3a*, enzymes that are expressed in the lens but not obviously related to chaperone activity. It is interesting to note that lens proteins are subject to relatively high levels of oxidative stress, which is one of the major sources of SSBs to which XRCC1-dependent repair is a response ^15^. Intriguingly, one of the protein partners of human XRCC1 is acyl peptide hydrolase/oxidized protein hydrolase, an enzyme implicated in the degradation of oxidized proteins ^16^. It is thus tempting to speculate that the upregulation of crystallins detected here is a response to the reduced ability of *xrcc1*-depleted fish to respond to protein oxidation.

## Methods

### Zebrafish husbandry

Experimental procedures were approved by the NICHD Animal Care and Use Committee (21-007). All methods were carried out in accordance with relevant guidelines and regulations. All methods are reported in accordance with ARRIVE guidelines (https://arriveguidelines.org). Zebrafish larvae were anesthetized and euthanized using buffered tricaine methanesulfonate. Zebrafish were of the Tüpfel long fin strain background, ordered from the Zebrafish International Resource Center. Larvae were raised on a 14/10 h light/dark cycle at 28°C at a maximum density of 10 larvae per 10 mL E3 medium supplemented with 1.5 mM HEPES pH 7.3. Behavior, brain imaging and RNA-seq were performed in larvae up to 7 dpf, before sex differentiation. Zebrafish lines used were *TgBAC(slc17a6b[vglut2a]:loxP-DsRed-loxP-GFP)nns14)*, previously converted to GFP by injection of Cre mRNA (*vGlut2a:GFP*) ^17^, *TgBAC(gad1b:lox-RFP-lox-GFP)nns26 (gad1b:RFP)* ^17^ and *Tg(Cau*.*Tuba1a:mCardinal)y516* (*tuba:mCar*) ^12^.

### Mutagenesis

We used ChopChop to select CRISPR targets in *xrcc1* and *parp1* and to design PCR primers for genotyping (Supp. Fig. 1 and 4) ^18^. For CRISPR, we pre-formed ribonucleoprotein complexes ^19^ but using tracrRNA, crRNA and Cas9 protein instead of single guide RNAs (Alt-R reagents, IDT), and injected each fertilized embryo at the one cell stage with 41 nmol guide-RNA duplex and 766 pg Cas9. To measure the efficiency of guide RNAs we performed CRISPRstat ^20^. Briefly, CRISPRstat involves amplifying the genomic DNA surrounding each target site using fluorescently tagged primers, then using capillary electrophoresis to assess the spectrum of product sizes. In control embryos, the PCR product is almost entirely present in a single peak, whereas efficient gRNAs generate such a high rate of insertion-deletions that the wildtype peak is extinguished and replaced with a range of product sizes. We then used a custom script to measure the area under each peak reported by the ABI machine and calculated the fraction of the total area occupied by the wildtype peak. Targets and corresponding PCR primers for *xrcc1* and *parp1* are in Supp. Fig. 1 and 4 respectively, with the sequence TGTAAAACGACGGCCAGT added in front of the forward primer and the sequence GTGTCTT added before the reverse primer. Product sizes in Supp. Fig. 1b and 4b do not include these tags. In all experiments, control embryos were taken from the same clutches as crispants, and injected only with tracrRNA and Cas9 (i.e., lacking a targeting guide RNA).

### Behavior

For behavioral tests, we recorded responses at 1000 frames per second with a high-speed camera (DRS Lightning RDT/1; DEL Imaging) and analyzed videos for responsiveness and movement kinematics using Flote software ^21^. We used the Dark Flash test to assess visual responsiveness ^22^. In this assay, larvae respond to a sudden loss of illumination by executing a specific ‘O-bend’ maneuver. After adapting 7 dpf larvae to an illumination level of 500 µW/cm^2^, we presented larvae with a pseudorandom sequence of reductions in light intensity to either 150 µW/cm^2^ (dim flash) or to 0 µW/cm^2^ (dark flash) and recorded the next 800 ms of movement with a high speed camera (1000 Hz). Stimuli were separated by 1 minute to avoid habituation. To assess auditory responsiveness and sensorimotor gating, we used a digital-to-analog card (PCI-6221; National Instruments) to generate 2 ms duration waveforms, delivered to larvae using a minishaker (4810, Bruel and Kjaer) ^23^. We tested larvae with a series of 80 vibrational stimuli of four types in a pseudorandom sequence, separated by 15s: weak, strong, weak with a 50 ms interval then strong, weak with a 500 ms interval then strong. The weak stimulus was 10% of the intensity of the strong stimulus, which had been calibrated so that larvae responded with a short-latency C-start on around 50% of trials. Responses to the weak and strong stimuli in isolation measured auditory responsiveness. Prepulse inhibition (at 50 and 500 ms) was measured using the double-stimulus trials. For habituation, we compared mean percent responsiveness to strong stimulus trials in the first half of the assay to mean responsiveness in the second half of the assay. As the camera was positioned above the arena, larvae with balance defects – lying on their sides – were detected as they only presented one eye to the camera ^24^.

### Brain morphometry

To perform brain morphometry, we crossed triple transgenic *vGlut2a*:*GFP, gad1b:RFP, tuba:mCar* fish to wildtypes, and injected embryos with CRISPR reagents. We sorted for embryos that were GFP+, RFP+ and mCar+, then scanned the brains of 10-20 larvae at 6 dpf using a confocal microscope (Nikon AI, 20x objective) ^25^. After scanning, we assessed CRISPR efficacy in each larva using CRISPRstat, and included crispants for further analysis if at least one guide RNA reduced the wildtype peak to less than 50% of the total PCR product. We registered each brain scan to the Zebrafish Brain Browser (ZBB), using multi-channel elastic registration with ANTs ^26^. We then used the resulting transformation matrix for each brain to inversely register the ZBB neuroanatomical annotations to the original brain scans, so that we could measure the volume of each brain and its divisions. We used the ANTS *CreateJacobianDeterminantImage* function on the transformation matrix for each larva to create a map of voxels that were dilated or compressed in volume compared to the reference brain (log Jacobian Determinant, LJD). Finally, we used Cobraz software to compare the fluorescence intensity and voxel volume in controls and mutants ^12^. To combine data from two crispant experiments, we used a custom Python script to calculate the ANOVA F-score for each voxel in brain maps of GFP, RFP, mCar and the LJD map to identify the main effects of genotype and experimental batch. For Figure 2E, we registered previously reported fluorescent in situ image stacks for *en1b, sox2, zic2a* to ZBB using an inter-atlas transformation matrix ^27,28^. In *parp1* rescue experiments, CRISPR reagents were injected into single transgenic *vGlut2a*:*GFP* embryos, as this pattern is sufficient to drive precise registration.

### Transcriptomics

To identify genes that were differentially expressed in both gRNA-t1/2 crispants and gRNA-t5/6 crispants, we extracted total RNA from larval heads using a Maxwell 16 LEV Simply RNA Tissue kit (Promega), then performed quality control using an Agilent Bioanalyzer. DNA extracted from tails was used to confirm CRISPR efficacy in each larva. Each biological replicate comprised RNA pooled from 2-3 larvae, and we used 3 replicates for each condition. The NICHD Molecular Genomics Core prepared and sequenced cDNA libraries. Quality assessment of the raw FASTQ files was conducted using FastQC v.0.12.1, followed by trimming with Cutadapt v.4.4 to remove adapters and low-quality sequences. Trimmed reads were then aligned to the zebrafish genome (GRCz11/danRer11) using STAR v.2.7, generating BAM files, from which we derived counts per gene using Subread v.2.0.3. We used DeSeq2 (v1.33.0) with apeglm shrinkage to identify genes that were differentially expressed between controls and crispants ^29,30^. Next, we filtered out genes with base mean expression less than 100 in either experiment, resulting in a total of 17,629 genes for further analysis. We then applied the Benjamini-Hochberg false discovery rate (FDR) correction to the p-values reported by DeSeq2 for the remaining genes. Genes with an FDR-adjusted p-value of less than 0.1 were considered significant.

### Statistics

We used the dabest Python library to analyze confidence intervals and generate estimation plots ^31^. ANOVAs were performed using JASP ^32^. T-tests are two-way independent sample tests. In boxplots, center line indicates median, box limits are quartiles and whiskers show furthest point within 1.5× interquartile range. For all tests, we used alpha 0.05, except for false discovery rate corrected p-values for differential gene expression which was 0.1.

## Data Availability Statement

Brain imaging data that support the findings of this study have been deposited at Zenodo and is available at: https://doi.org/10.5281/zenodo.14036962. RNA sequencing data is available at the NCBI Sequence Read Archive repository (https://dataview.ncbi.nlm.nih.gov/object/PRJNA1188175).

## Acknowledgments

H.B. discloses support by the Intramural Research Program of the Eunice Kennedy Shriver National Institute for Child Health and Human Development (NICHD). This study utilized the high-performance computational capabilities of the Biowulf Linux cluster at the National Institutes of Health, Bethesda, MD. K.W.C. discloses support by program grants from the MRC (MR/W024128/1) and CRUK (C6563/A27322). R.M. discloses support from international collaborators biobanks, and for essential funding from The Wellcome Trust, The MRC, The MSA Trust, The National Institute for Health Research University College London Hospitals Biomedical Research Centre NIHR-BRC), The Michael J Fox Foundation (MJFF), The Fidelity Trust, Rosetrees Trust, The Dolby Family fund, Alzheimer’s Research UK (ARUK), MSA Coalition, Parkinson’s disease society, Parkinson’s Foundation, The Guarantors of Brain, Cerebral Palsy Alliance, FARA, EAN, Victoria Brain bank, the NIH NeuroBioBank, Queen Square BrainBank, The MRC Brainbank Network.

