## Supplemental Material for "*Parp1* deletion rescues cerebellar hypotrophy in *xrcc1* mutant zebrafish"

**A**

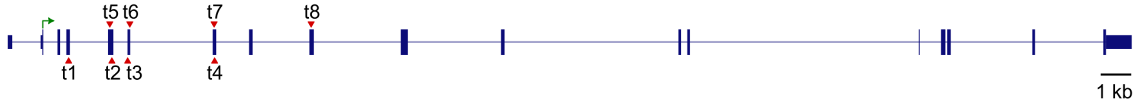


**B**

| gRNA | set | exon | target sequence | genotyping primer F | genotyping primer R | size |
|---|---|---|---|---|---|---|
| t1 | 1 | 4 | CAGACGTGGAGTGTCCGACAAGG | TCCTCCAGTTGGAAAAGGAAGAG | AACTGACTGAAGACAAAACTCACT | 197 |
| t2 | 1 | 5 | GCTGGTGAAGGCGCAATCACAGG | GCATTGTTTTGTAGGTTCTGCTGG | CCCCTGACCTTGCTGTAGGG | 181 |
| t3 | 1 | 6 | TGTGTGCAGACTATAGCGTACGG | ATGTTTAACCACACCCTTCCAG | TTTAGATGCAAAACACTCACCG | 214 |
| t4 | 1 | 7 | AAAGACGAGTCTCCTTCTGCTGG | GTTCTTCACTTTGAAAGCACCC | GGATTTAGCTGAAGTGTCCCTG | 225 |
| t5 | 2 | 5 | CGTGTGCGTTTCTTCGGGCCCGG | CCGCATGTTTACTCTTCTGCAT | TTCACGCGATCCCACTTCTC | 163 |
| t6 | 2 | 6 | TCACCGGTGGAGTCGAGACTAGG | TATAGCGTACGGCATCTCCTTC | ACACTTTCCGGATGCTCTGTAT | 243 |
| t7 | 2 | 7 | GCTGGGTCAAACGTGCAGCCTGG | ACATCTGTCTGCTCTCTCCCTC | ACTACGGTGCTTTACCCTTCAG | 169 |
| t8 | 2 | 9 | TCGGCAGATGCAGATGAACTGGG | TGAAGCGAAAGTTTGAGTTCAG | TTCATTTGTACCTCTGCTGCTG | 152 |

**C**

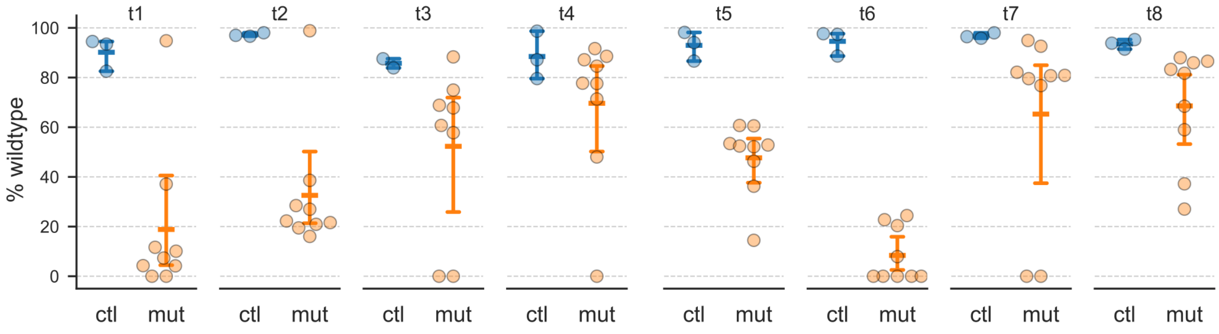


**Supplemental Figure 1. Active guide RNAs targeting *xrcc1***

A. Location of guide RNAs tested for efficacy against *xrcc1* exons. The first set of gRNAs (t1-4) were co-injected in the first experiment, and the second set (gRNAs t5-8) co-injected for the replication experiment. B. gRNA target sequence, genotyping primers and PCR product size (base pairs). C. Proportion of PCR product present in the wildtype sized ABI peak in individual embryos for controls that were injected with Cas9 protein and tracR RNA only (blue) or Cas9 protein, tracR RNA and targeting gRNA (orange).

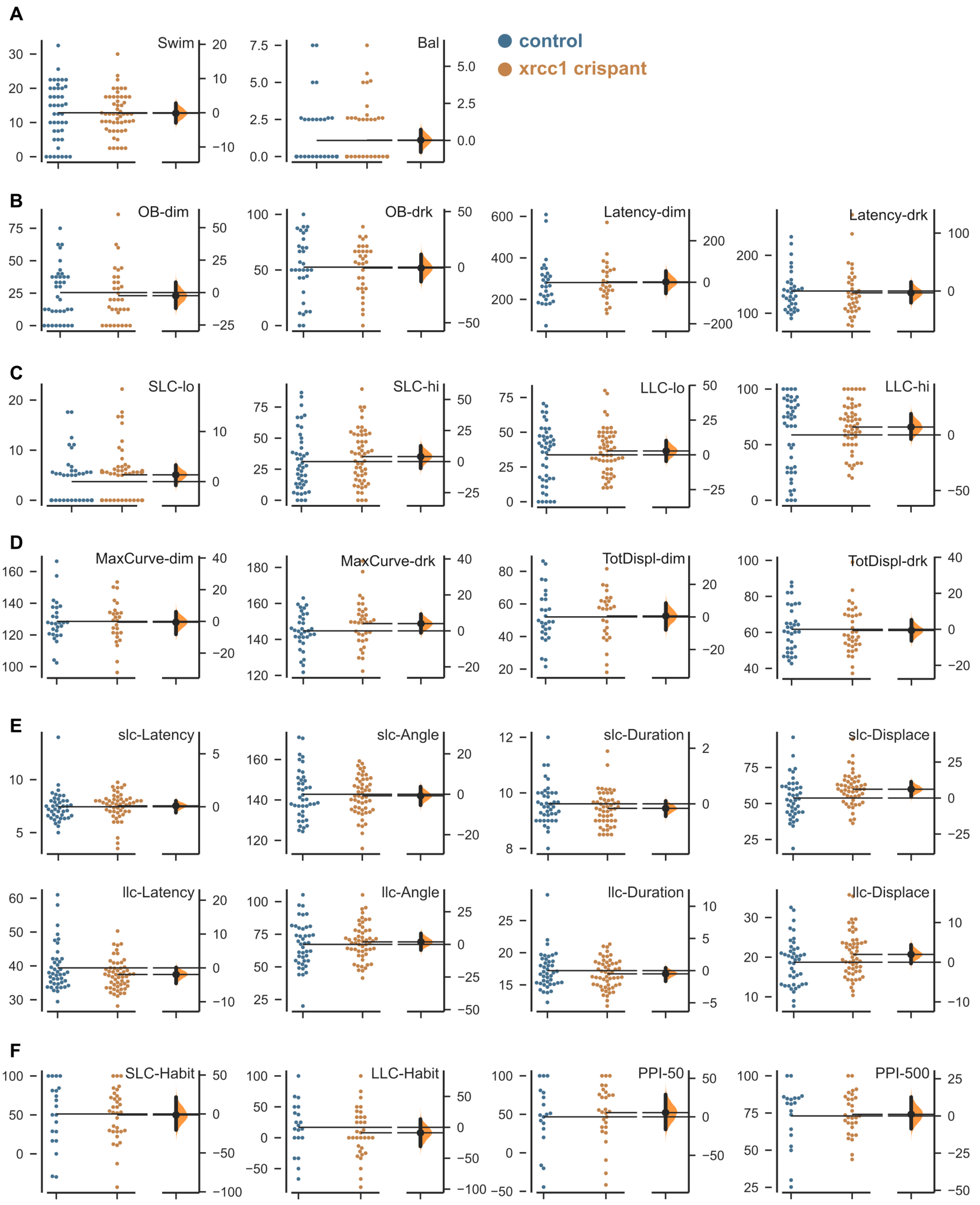

A
B
C
D
E
F
control
xrcc1 crispant
Swim
Bal
OB-dim
OB-drk
Latency-dim
Latency-drk
SLC-lo
SLC-hi
LLC-lo
LLC-hi
MaxCurve-dim
MaxCurve-drk
TotDispl-dim
TotDispl-drk
slc-Latency
slc-Angle
slc-Duration
slc-Displace
llc-Latency
llc-Angle
llc-Duration
llc-Displace
SLC-Habit
LLC-Habit
PPI-50
PPI-500

**Supplemental Figure 2. Sensory and motor behavior in *xrcc1* crispants**

Estimator plots showing measurements of sensory, motor and central integration in controls (blue) and *xrcc1* crispants (orange). Tests measure spontaneous swimming and balance (A), visual sensory responsiveness (B), auditory sensory responsiveness (C), movement kinematics after visual cue (D), movement kinematics after auditory cue (E) and sensorimotor processing (F). Estimator plots show mean measurements for each larva tested on the left side, and mean difference between controls and crispants on right. The slc-Displace difference was nominally significantly different (p=0.023), but not after adjustment for multiple comparisons. See supplemental table 1 for explanation of measurements. N=18 to 54 larvae per group.

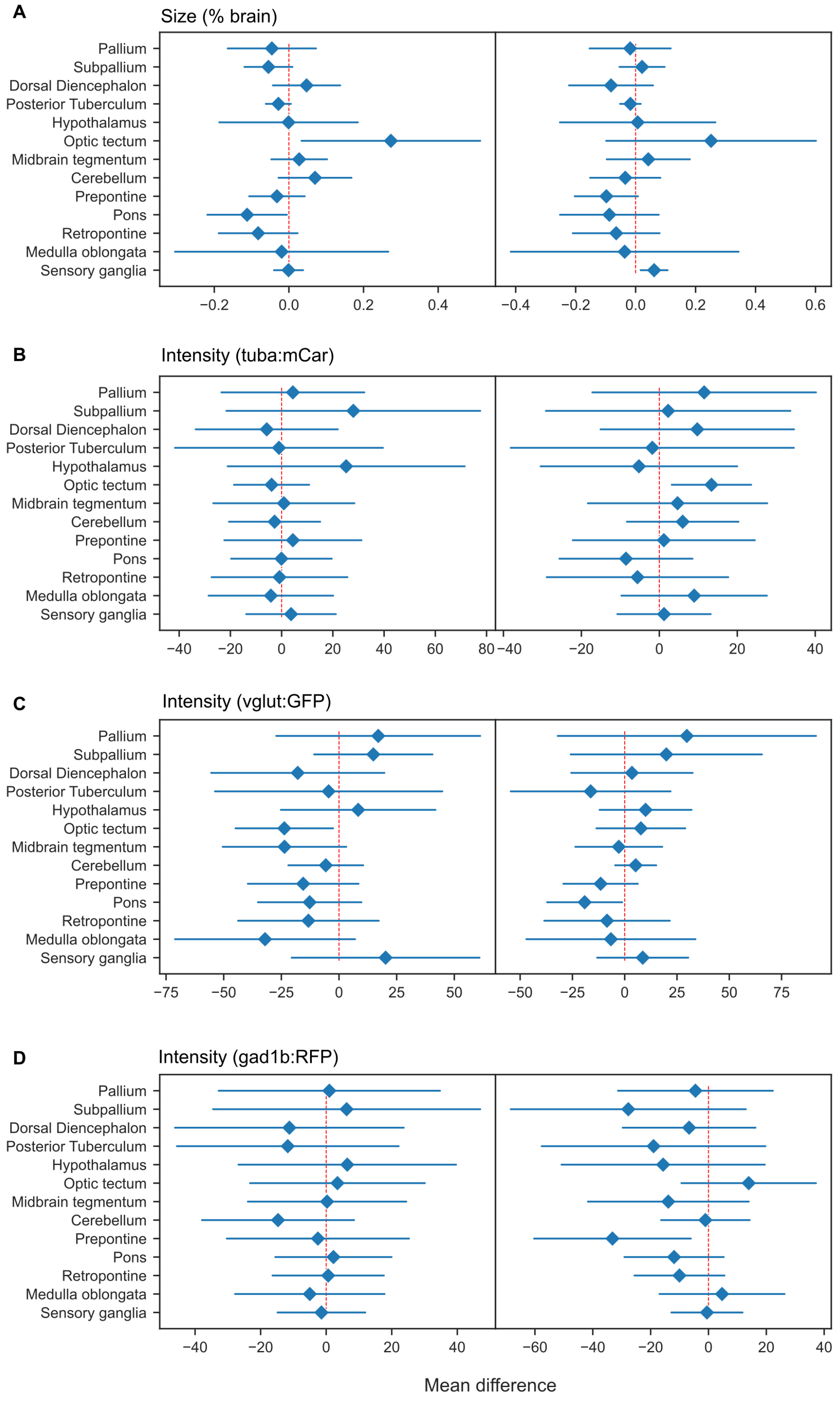

A
Size (% brain)
B
Intensity (tuba:mCar)
C
Intensity (vglut:GFP)
D
Intensity (gad1b:RFP)
Pallium
Subpallium
Dorsal Diencephalon
Posterior Tuberculum
Hypothalamus
Optic tectum
Midbrain tegmentum
Cerebellum
Prepontine
Pons
Retropontine
Medulla oblongata
Sensory ganglia
Mean difference

**Supplemental Figure 3. Brain morphometric measurements in *xrcc1* crispants**

Mean difference and confidence interval for size (A) and fluorescence intensity (B-D) in brain divisions in *xrcc1* crispants. Division size was normalized as the percentage of total brain volume. Fluorescence intensity is in arbitrary units. Left panels show data for gRNA-t1/2 crispants, right panels for gRNA-t5/6. Mean difference values greater than zero indicate a larger size or greater intensity in controls.

**A**

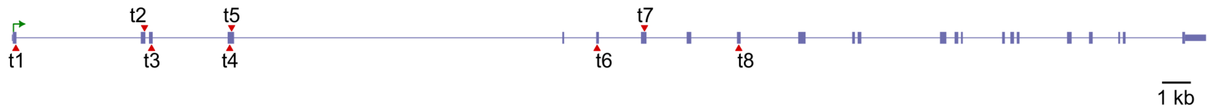


**B**

| gRNA | set | exon | target sequence | genotyping primer F | genotyping primer R | size |
|---|---|---|---|---|---|---|
| t1 | 1 | 1 | TGCTAAGGACTCGCTGAGGA | CAGGACGACAAGCTGTACAAAG | CCAAAAAGTTGCACACAACAAT | 144 |
| t2 | 2 | 2 | GTGAAGACAGCCATTGAGAG | ATTTACTGACCTCCGCTGGG | AAGACTAACTGCTCACCAGCC | 150 |
| t3 | 2 | 3 | TTTCTGATCGCAGCCCTTGC | ATCCTTGTTGCTTTACAGGGAA | AGCATTGGGCCTAAGTAATTCA | 207 |
| t4 | 1 | 4 | CCGGCTGCTTTGTGAGTCGT | TTTTTGTTGTTGTTCCCTGATG | CTCGTAGAACTGCGAATCCTTT | 193 |
| t5 | 2 | 4 | AGACCAAGCTGAGGTTTCTC | TTTTTGTTGTTGTTCCCTGATG | CTCGTAGAACTGCGAATCCTTT | 193 |
| t6 | 1 | 6 | CCAGCTGATCTGGGGAATTA | CCTTTCTTCCTCATTCTGATGC | GCATTCTTACATTTGACTCCCC | 273 |
| t7 | 2 | 7 | AAGCCTTGTGAGACTTGCAA | TGTGGATCGACTATCAGACTGC | CCGCTTGAAGGAGAGAAAATAG | 267 |
| t8 | 1 | 9 | TACTGAGCTTTCCCACAGCC | TACTTGGCTAAACTGACTGCGA | GATTTTTCCTCCCAGATCTTCC | 284 |

**C**

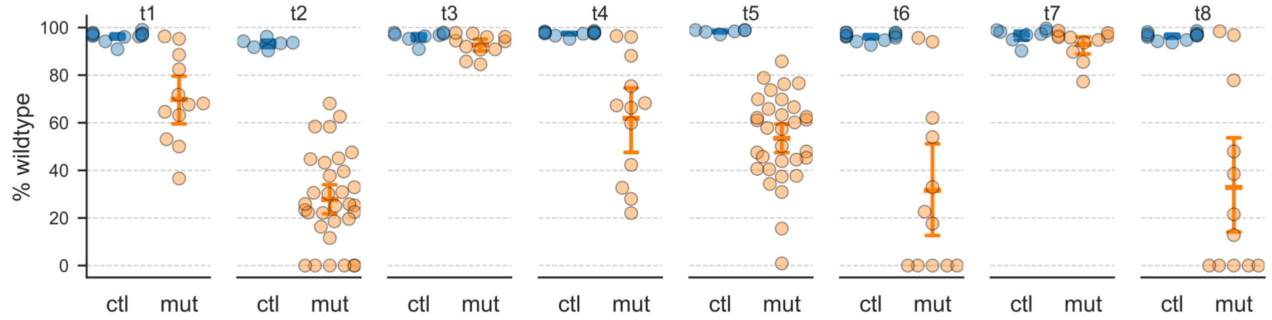


**Supplemental Figure 4. Active guide RNAs targeting *parp1***

A. Location of guide RNAs against *parp1* exons tested for cutting efficiency. B. Target sequence and PCR primers for CRISPRstat genotyping for *parp1* guide RNAs. C. Cutting efficiency for *parp1* guide RNAs, shown as the percentage of the normal wildtype peak remaining after injection of each guide RNA during CRISPR experiments.

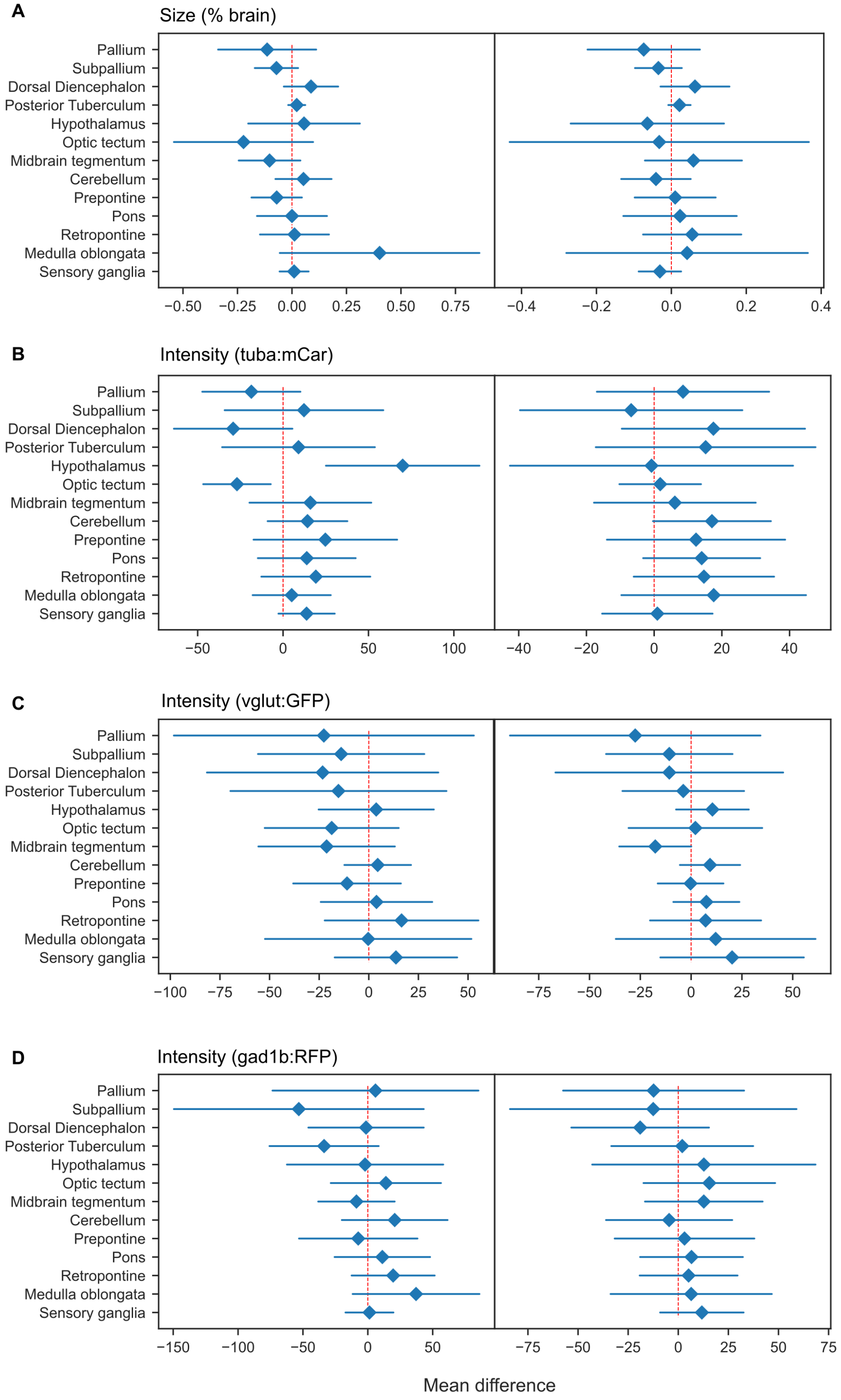

A
Size (% brain)
B
Intensity (tuba:mCar)
C
Intensity (vglut:GFP)
D
Intensity (gad1b:RFP)
Pallium
Subpallium
Dorsal Diencephalon
Posterior Tuberculum
Hypothalamus
Optic tectum
Midbrain tegmentum
Cerebellum
Prepontine
Pons
Retropontine
Medulla oblongata
Sensory ganglia
Mean difference

**Supplemental Figure 5. Brain morphometric measurements in *parp1* crispants.**

Mean difference and confidence interval for size (A) and fluorescence intensity (B-D) in brain divisions in 6 dpf *parp1* crispants. Left panels: gRNA-t2/5. Right panels: gRNA-t6/8.

| Label | Meaning | Measurement |
|---|---|---|
| Swim | Spontaneous swimming activity | Percent larvae active in 30 ms window before presentation of auditory cue |
| Bal | Balance | Percent larvae failing to present with dorsal side up |
| OB-dim | Visual responsiveness (dim flash) | Percent larvae responding to small light decrement with O-bend |
| OB-drk | Visual responsiveness (dark flash) | Percent larvae responding to light extinction with O-bend |
| Latency-dim | Visual responsiveness (dim flash) | Response latency (ms) for larvae with O-bend response to small light decrement |
| Latency-drk | Visual responsiveness (dark flash) | Response latency (ms) for larvae with O-bend response to light extinction |
| SLC-lo | Auditory responsiveness (weak auditory cue) | Percent larvae responding to weak auditory cue with short latency C-start |
| SLC-hi | Auditory responsiveness (intense auditory cue) | Percent larvae responding to intense auditory cue with short latency C-start |
| LLC-lo | Auditory responsiveness (weak auditory cue) | Percent larvae responding to weak auditory cue with long latency C-start |
| LLC-hi | Auditory responsiveness (intense auditory cue) | Percent larvae responding to intense auditory cue with long latency C-start |
| MaxCurve-dim | Movement kinematics (dim flash) | Mean maximal curvature (degrees) during O-bend responses to small light decrement |
| MaxCurve-drk | Movement kinematics (dark flash) | Mean maximal curvature (degrees) during O-bend responses to light extinction |
| TotDispl-dim | Movement kinematics (dim flash) | Mean net displacement (pixels) during O-bend responses to small light decrement |
| TotDispl-drk | Movement kinematics (dark flash) | Mean net displacement (pixels) during O-bend responses to light extinction |
| slc-Latency | Movement kinematics (short latency c-starts) | Mean latency (ms) to initiation of short latency C-start in response to auditory cue |
| slc-Angle | Movement kinematics (short latency c-starts) | Mean maximal change in head orientation (degrees) during initial bend of short latency C-start to auditory cue |
| slc-Duration | Movement kinematics (short latency c-starts) | Mean time (ms) until maximal change in head orientation during initial bend of short latency C-start to auditory cue |
| slc-Displace | Movement kinematics (short latency c-starts) | Mean net displacement (pixels) during short latency C-start to auditory cue |
| llc-Latency | Movement kinematics (long latency c-starts) | Mean latency (ms) to initiation of long latency C-start in response to auditory cue |
| llc-Angle | Movement kinematics (long latency c-starts) | Mean maximal change in head orientation (degrees) during initial bend of long latency C-start to auditory cue |
| llc-Duration | Movement kinematics (long latency c-starts) | Mean time (ms) until maximal change in head orientation during initial bend of long latency C-start to auditory cue |
| llc-Displace | Movement kinematics (long latency c-starts) | Mean net displacement (pixels) during long latency C-start to auditory cue |
| SLC-Habit | Sensorimotor filtering | Percent habituation of short latency C-start responses across trials conducted with intense auditory cue |
| LLC-Habit | Sensorimotor filtering | Percent habituation of long latency C-start responses across trials |

| | | |
|---|---|---|
| | | conducted with intense auditory cue |
| PPI-50 | Sensorimotor filtering | Percent prepulse inhibition of short latency C-starts when intense auditory cue preceded at 50 ms by weak auditory prepulse |
| PPI-500 | Sensorimotor filtering | Percent prepulse inhibition of short latency C-starts when intense auditory cue preceded at 500 ms by weak auditory prepulse |

**Supplemental Table 1**

Definitions of abbreviations for measurements of sensory and motor measurements in supplemental figure 2.
